# *In vivo* motor unit decoding and *in vitro* cellular characterisation of spinal circuits for urination in adult mice

**DOI:** 10.64898/2026.03.13.711618

**Authors:** MG Özyurt, F Nascimento, A Pascual-Valdunciel, K Dhillon, V Bansal, RM Brownstone, M Beato

**Author notes:** Equal Contribution (F Nascimento, A Pascual-Valdunciel).

## Abstract

Urinary dysfunction affects billions of individuals worldwide; however, the fundamental cellular and circuit properties that govern perineal motor control remain largely unknown, serving as a functional “black box”. Here, we describe several methods that, when used in concert, characterise cellular, synaptic, and motor unit properties underlying the control of urination in adult mice. High-density electromyography combined with real-time cystometry were used to study external urethral sphincter (EUS) motor units, which follow a hierarchical (“onion skin”) recruitment pattern during bladder filling. The transition to the voiding phase is marked by inhibition, followed by synchronised bursts. Furthermore, through concurrent recordings of ischiocavernosus (IC) muscles, the relationship between IC and EUS motor units could be studied to look for shared common inputs that could shed light on circuitry. Whole-cell patch-clamp recordings from retrogradely identified neurons revealed a fundamental biophysical divergence: urinary parasympathetic preganglionic neurons (PPGN) are significantly smaller and more excitable than somatic EUS and IC motoneurons and lack the recurrent excitatory and inhibitory circuits present in both EUS and IC motor pools. Finally, using a novel pressure-clamp preparation, we showed that acute tibial nerve stimulation (a widely used treatment for urinary dysfunction) evokes short-latency inhibition of EUS motor units. Collectively, these methods can be used to delineate patterns of motor unit recruitment, local recurrent microcircuit architecture, and distinct biophysical properties of the perineal motor system, providing mechanistic insights into urinary function.

## Introduction

Urination is a fundamental behaviour essential for maintaining fluid balance. Dysfunctions of the urinary system affect an estimated two to four billion adults globally ^(1–3)^, with urinary incontinence alone impacting over 400 million individuals ^(1)^. This condition diminishes personal autonomy and imposes significant demands on caregivers and healthcare systems, resulting in an economic burden of approximately €80 billion annually within the European Union alone ^(3–5)^. Despite this substantial impact, our poor understanding of the neural circuits that coordinate urination limits the development of targeted therapeutic interventions to mitigate this burden.

Neural control of urination operates hierarchically and is a product of a complex integration of autonomic and somatic nervous systems ^(6)^. While supraspinal control centres, such as the medial prefrontal cortex and the pontine micturition centre (also known as Barrington’s nucleus), are responsible for decision-making and initiation of the urination command ^(7–10)^, the final execution of this behaviour relies on microcircuits located in the lumbosacral spinal cord ^(6)^. Spinal neurons controlling the external and internal urethral sphincters (EUS and IUS, respectively) and associated perineal muscles must operate as a tightly coordinated ensemble, sustaining tonic activity during storage yet rapidly reconfiguring their output to permit efficient voiding ^(11)^. This coordination depends on local spinal networks that transform descending supraspinal commands and local sensory feedback into precise motor outputs.

Despite the criticality of lumbosacral spinal circuits in micturition, their specific cellular and network properties remain largely unknown. We know from studying other systems that opening this “black box” means determining the fundamental properties of its component neurons, their connectivity, and their activity during the behaviour in question. Characterising the cellular and circuit properties of perineal neurons and their recruitment pattern during urination is a critical step toward improving the development of precise, mechanism-driven neuromodulation and therapeutic strategies.

A prominent example of this translational gap is percutaneous tibial nerve stimulation, a widely used clinical neuromodulation therapy for overactive bladder and urinary incontinence ^(12,13)^. While chronic stimulation of the distal tibial nerve can alleviate some lower urinary tract symptoms, its underlying physiological mechanism of action remains largely speculative, often presumed to rely on the general modulation of sacral spinal reflex pathways ^(14,15)^. Importantly, relying on empirical clinical success without mechanistic insight is a significant barrier to developing next-generation, precision-targeted neuromodulatory devices and/or other therapies. Without a foundational understanding of the local lumbosacral microcircuits, it is not possible to determine how this peripheral input influences the specific perineal motor pools that govern urination.

Functionally characterising the spinal circuits governing urination requires robust characterisation of the high-resolution activity of the target muscles from storage to voiding, and experimental access to the cellular properties of the network elements. To achieve this, we developed a multi-scale methodological approach targeting EUS and ischiocavernosus (IC) muscles, whose motoneurons are intermingled within the Onuf’s nucleus, as well as urinary sphincter-related parasympathetic preganglionic neurons (PPGN). In adult mice, we used intramuscular high-density EMG recordings (HDEMG, using myomatrix arrays ^(16)^) from the EUS and IC during voiding and decomposed the activity of individual motor units to reveal their recruitment and firing patterns. To study intrinsic cellular and circuit properties, we retrogradely labelled the neurons innervating these muscles and characterised their intrinsic properties and recurrent synaptic inputs using whole-cell patch-clamp recordings in adult *in vitro* spinal cord preparations. Finally, we developed an *in vivo* ‘pressure-clamp’ preparation to establish controlled bladder distension, enabling the recording of continuous and stable EUS motor unit discharges. We used this approach to test the effect of tibial nerve stimulation on EUS motor unit firing. Together, these methods provide a comprehensive experimental framework for interrogating the spinal orchestration of urination and the physiological mechanisms of neuromodulation.

## Results

### Gradual recruitment of external urethral sphincter motor units and rate coding characterise the guarding and voiding reflexes

Unlike conventional bipolar EMG, which records global muscle activity lacking spatial precision, HDEMG captures the spatiotemporal distribution of motor unit potentials, allowing for the investigation of neural drive at the single neuron level. We used multichannel intramuscular Myomatrix arrays ^(16)^ to record from EUS and isolate individual motor unit activity throughout urinary cycles in four mice (three male, one female, total of 20 motor units) (**Figure 1A**). In urethane-anaesthetised animals, we artificially controlled bladder pressure with a constant flow of saline (25 µL/min ^(17)^) to examine sphincter motor unit activity in response to gradual increases in bladder pressure (guarding reflex), and during detrusor muscle contraction and sphincter relaxation during emptying (voiding reflex). During urination, the shift from storage to voiding requires a change in neural drive to spinal circuits, which in turn directly affects their activation pattern ^(6,18)^. We tracked individual units to answer a specific physiological question: is the continence/guarding reflex achieved by recruiting entirely new populations of motor units, or simply by a larger drive to an increased discharge of already active motor units, or both?

**Figure 1.**
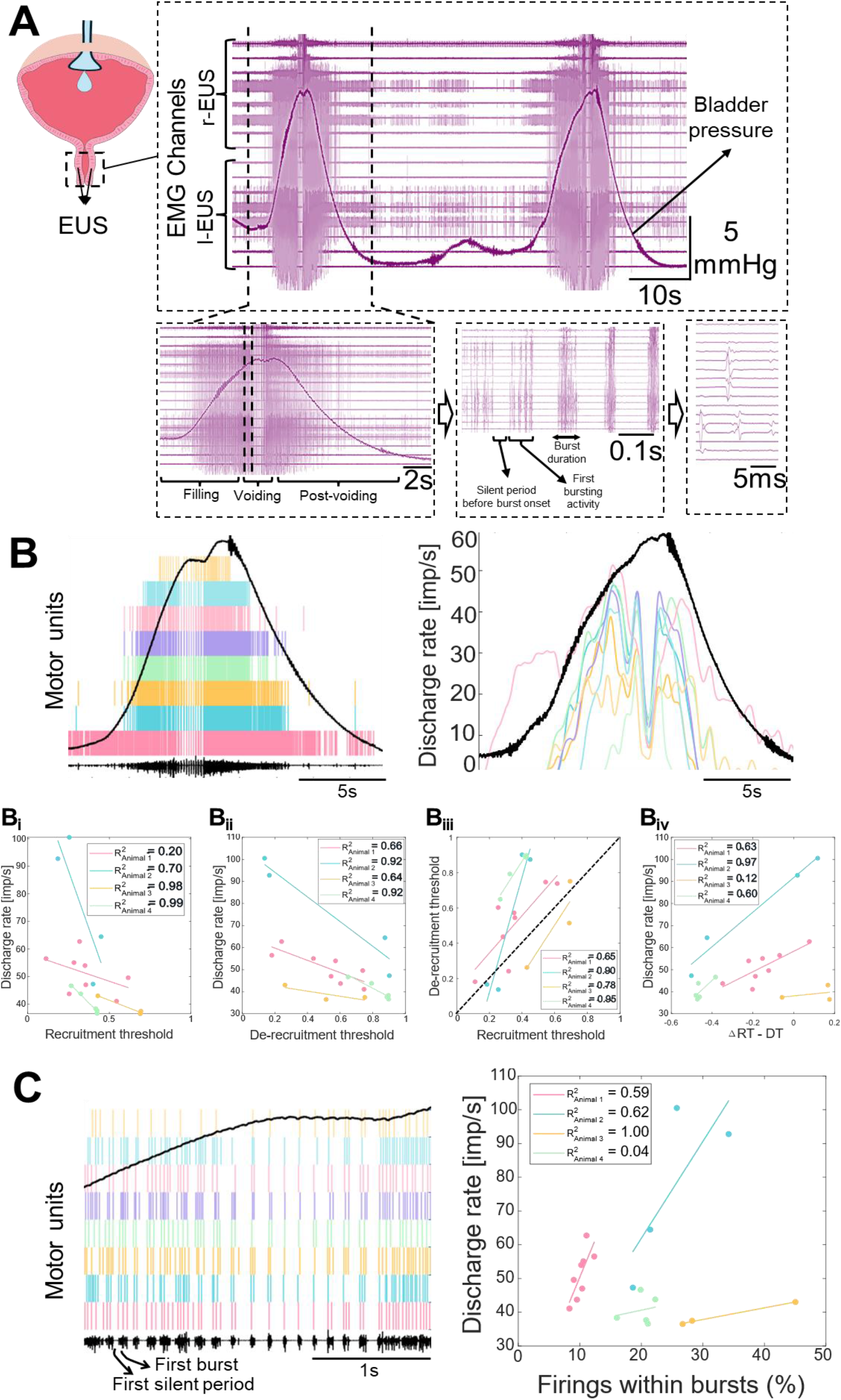
High-density EMG characterisation of external urethral sphincter (EUS) activity during cystometry. **(A)** Representative HDEMG recordings of EUS motor unit activity across two complete urinary cycles, illustrating the filling (storage) and voiding phases. **(B)** Analysis of motor unit firing properties, (**B**_**i**_**-B**_**iv**_) showing recruitment and de-recruitment discharge rates plotted as a function of bladder pressure (black line), where the threshold level 1 is the peak bladder pressure. The dashed diagonal line in B_iii_ indicates the line of unity. Data points falling below this line represent motor units that exhibit persistent firing. **(C)** Quantification of burst characteristics observed during the voiding phase. Data compiled from n = 20 motor units recorded in N = 4 mice (3 males, 1 female).

Motor unit analysis revealed the hierarchical strategy in maintaining the “guarding reflex” (**Figure 1B**). As bladder pressure rises, we observed a systematic increase in neural drive to EUS motoneurons, recruiting new motor units gradually while simultaneously increasing the discharge rate of those already active. This behaviour mirrors the “onion skin” phenomenon commonly seen in other skeletal muscles, where early-recruited units sustain higher firing rates than later-recruited ones ^(19–21)^. This suggests that the highly specialised EUS muscle shares fundamental control principles with the rest of the skeletal motor system.

In detailing these recruitment dynamics, we observed a linear relationship between the discharge rate and the recruitment threshold (with respect to bladder pressure) of EUS motor units (**Figure 1Bi**). This linear relationship is conserved during the de-recruitment phase (**Figure 1Bii**). However, the de-recruitment profile exhibits a pronounced offset; most of these units systematically stop discharging at a higher bladder pressure than the threshold at which they were initially recruited (**Figure 1Biii**). Notably, the magnitude of this hysteresis is not uniform across the motor pool; rather, it is larger for the later-recruited motor units, indicating that these units are de-recruited earlier as bladder pressure declines. (**Figure 1Biv**).

Finally, we focused on the bursting activity that marks the transition from the guarding to voiding reflex (**Table 1**). There is a notable species difference: while this transition in humans is characterised by the complete silence of motor units, in mice it is marked by distinct bursting activity ^(18,22)^. Interestingly, these bursts (at ~6.5 Hz rate and each last around 47 ms, see **Table 1**) gradually synchronise before the peak bladder pressure. The initial burst is usually preceded by a robust inhibition (**Figure 1C**). These bursts contain a progressively higher number of discharges in the earlier-recruited motor units (i.e., higher discharge rate units) (**Figure 1C** – right panel), indicating that the hierarchical recruitment pattern is also preserved within these transient bursts.

**Table 1.** Burst properties during voiding phase.

| Burst frequency (Hz) | Burst duration (ms) | Burst start (normalised to max bladder pressure) | Burst end (normalised to max bladder pressure) |
| --- | --- | --- | --- |
| 6.5 [7.4 5.6] | 47 [49 45] | 0.70 [0.87 0.41] | 0.80 [0.91 0.70] |

*Median [95% CI]*

Collectively, these findings suggest that the guarding phase is defined by the recruitment of new motor units combined with an increase in firing rates relative to rising bladder pressure, which could indicate that there is a common input homogeneously projected to all motor units. Different intrinsic properties across motoneurons could explain the gradual motor unit recruitment and de-recruitment.

### The ischiocavernosus muscle is active in both the guarding and voiding phases of the urination cycle

Although the EUS is the primary somatic muscle responsible for urination, it functions within a broader perineal muscle synergy. Evidence suggests that the IC motor units are also recruited during urination in humans ^(23)^. Given this clinical evidence, and because pelvic reflexes frequently recruit perineal muscles as integrated functional units rather than isolated targets ^(24)^, we aimed to understand the specific activation pattern of the IC during micturition.

We did so through HDEMG recordings of the IC muscle and found that the IC exhibits an activation pattern temporally correlated with the EUS during both guarding and voiding (**Figure 2A**). To determine whether this synchronisation reflects a common neural drive rather than independent recruitment, we quantified intramuscular coherence, a method that quantifies the degree of correlated oscillatory activity between signals in the frequency domain. We investigated coherence using motor units or rectified EMG activity within EUS muscle (intramuscular, three male and one female mice) (**Figure 2B**) as well as between ipsilateral EUS and IC muscle (intermuscular) during cystometry in a male mouse (example shown in Figure 2C). Interestingly, the results indicate a coherence peak at ~8 Hz both within the EUS and across the EUS-IC motor pools. This suggests that these anatomically and functionally distinct muscles receive shared local or supraspinal inputs during guarding and voiding, supporting the hypothesis that the IC may play a role in urination.

**Figure 2.**
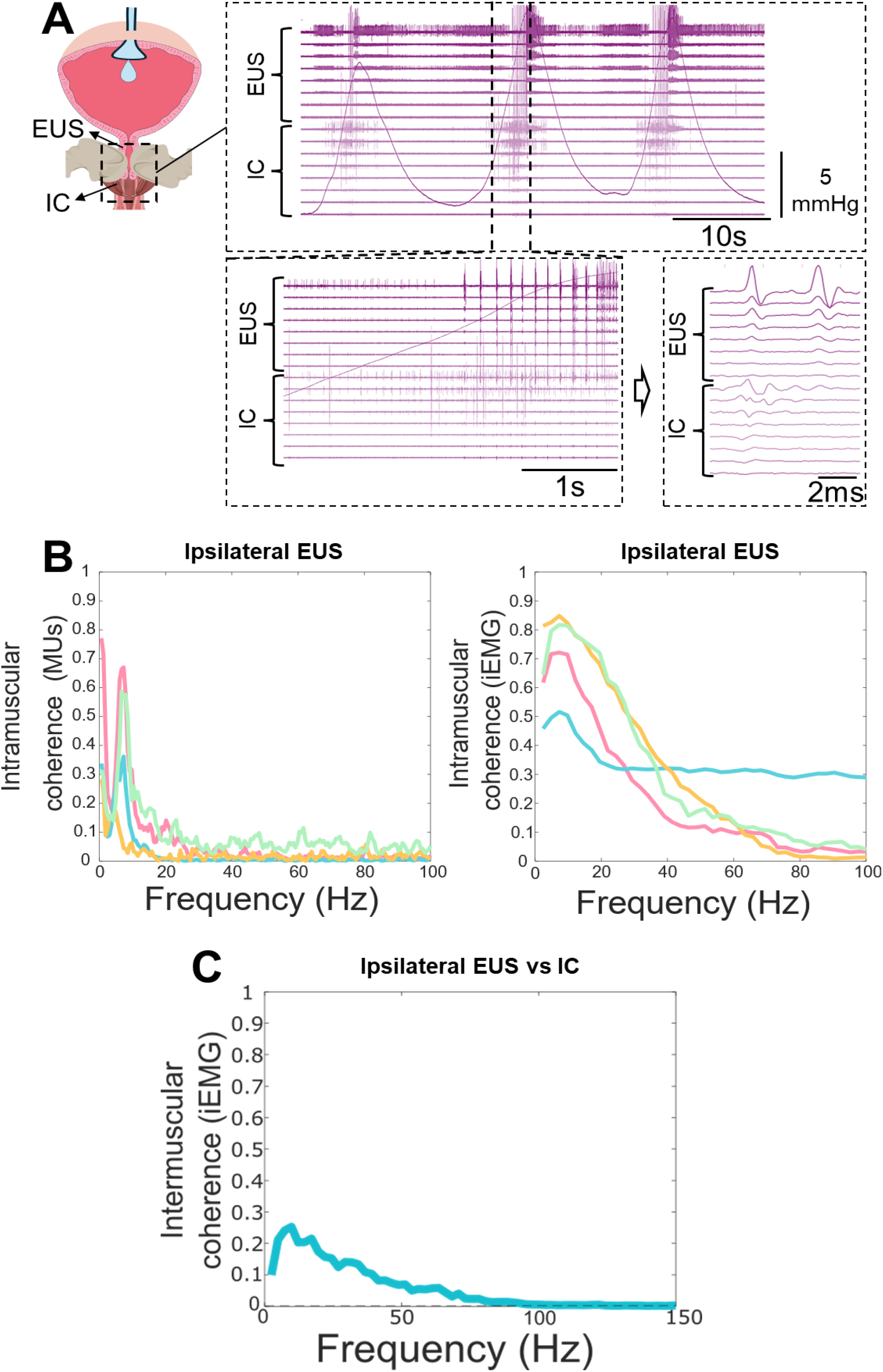
High-density EMG characterisation of ischiocavernosus (IC) activity during cystometry. **(A)** Representative HDEMG recordings of IC and external urethral sphincter, (EUS) motor unit activity across three complete urinary cycles and zoomed panels. **(B)** Analysis of intramuscular coherence using motor units (MUs, left) and integrated EMG (iEMG, right) within EUS muscle (3 male, 1 female mice, each coloured line is an animal). **(C)** Intermuscular coherence value between ipsilateral EUS and IC muscle activation using iEMG (a male mouse).

### Somatic motoneurons and parasympathetic preganglionic neurons exhibit divergent cellular properties

While the EUS and IC represent some of the somatic components of pelvic floor control, the PPGN is the critical autonomic counterpart. To determine if they share similar cellular properties, that could give rise to similar firing patterns or excitability profile, we performed whole-cell patch-clamp recordings from retrogradely labelled PPGN and compared their intrinsic properties to the somatic EUS and IC motor pools (**Figure 3A and 3B**).

**Figure 3.**
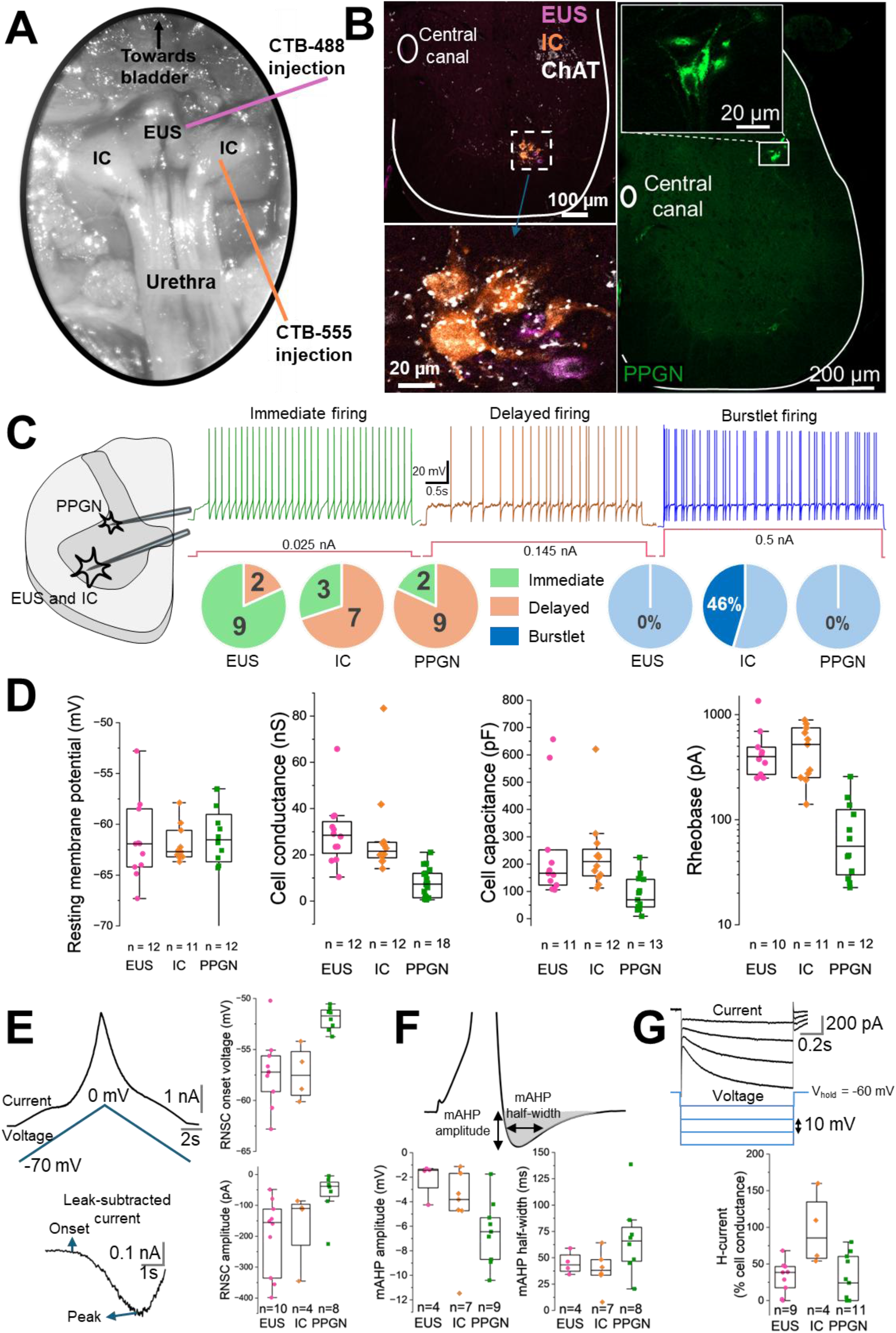
Active and passive properties of somatic and autonomic perineal neurons. **(A)** Image illustrating the retrograde tracer injection sites in the urethral sphincter and ischiocavernosus (IC) muscles. **(B)** Representative image of the L6-S1 spinal cord showing retrogradely labelled (with cholera toxin subunit β, CTB) external urethral sphincter (EUS) and IC motoneurons alongside parasympathetic preganglionic neurons (PPGN). **(C)** Representative drawing and current-clamp traces displaying the distinct firing phenotypes of these neuronal populations. **(D)** Quantification of basic membrane properties; resting membrane potential, capacitance, conductance, rheobase. **(E– G)** Comparative analysis of active properties, including region of negative slope conductance (RNSC), medium afterhyperpolarisation (mAHP) and hyperpolarisation-activated current (H-current). Number of neurons (n) recorded from 11 mice (10 males, 1 female; 42–111 days old).

Our recordings revealed a clear biophysical distinction in firing phenotypes (**Figure 3C**). While limb motoneurons are classically categorised by their immediate or delayed firing in response to constant current injection ^(25)^, it remains unknown whether these putative ‘slow’ and ‘fast’ phenotypes applies to pelvic motoneurons. Interestingly, the majority of PPGN and IC motoneurons exhibited a delayed firing phenotype (9/11 for PPGN, 7/10 for IC), whereas most of EUS motoneurons showed immediate firing type (9/11). Moreover, we have observed a heterogeneous discharge profile within the IC motoneuron. Approximately half of the recorded IC motoneurons (46%) responded to constant current injection with a burstlet firing pattern. In contrast, this type of discharge was never observed in EUS or PPGN motoneurons. This suggests that a subset of IC motoneurons has specialised membrane properties capable of supporting the rhythmic, high-frequency contractions, reflecting a potential diversity of functions within this specific motor pool.

All neuron types exhibited similar resting membrane potentials at approximately −62 mV. However, distinct differences emerged in passive membrane properties. PPGN neurons systematically displayed significantly lower whole-cell conductance and capacitance (approximately 60% lower than somatic motoneurons) and a dramatically lower rheobase (approximately 90% lower than somatic motoneurons). The lower capacitance reflects a smaller soma surface area, while lower rheobase indicates higher excitability, allowing these autonomic neurons to be recruited by weaker synaptic inputs compared to their somatic counterparts (see **Table 2** for details and **Figure 3D**).

**Table 2.** Basic properties of perineal neurons.

|  | **RMP (mV)** | **Conductance (nS)** | **Capacitance (pF)** | **Rheobase (pA)** |
| --- | --- | --- | --- | --- |
| **EUS** | -62 [-59 -64] | 29 [38 21] | 167 [374 114] | 398 [724 252] |
| **IC** | -63 [-61 -63] | 22 [40 16] | 209 [321 149] | 520 [671 316] |
| **PPGN** | -62 [-59 -64] | 7 [11 5] | 69 [128 51] | 56 [130 39] |

*EUS: External urethral sphincter, IC: Ischiocavernosus, PPGN: Parasympathetic preganglionic neuron, RMP: resting membrane potential. Median [95% CI]*

In addition to differences in basic properties, these neurons display distinct active properties. While all recorded neurons exhibited Type 2 adaptive-type firing hysteresis, PPGN showed more depolarised region of negative slope conductance (RNSC) onset voltage and smaller RNSC amplitude compared to somatic motoneurons (**Figure 3E**). Similarly, PPGN neurons exhibited distinct mAHP properties, characterised by a larger amplitude and slightly longer half-width (**Figure 3F**). Interestingly, IC motoneurons displayed uniquely large H-current, with an amplitude approximately double that observed in PPGN and EUS motoneurons (**Table 3 and Figure 3G**).

**Table 3.** Active properties of perineal neurons.

|  | RNSC onset (mV) | RNSC amplitude (pA) | mAHP amplitude (mV) | mAHP half-width (ms) | H-current size (% cell conductance) |
| --- | --- | --- | --- | --- | --- |
| **EUS** | -57 [-55 -60] | -156 [-111 -286] | -1.4 [0.2 -4.4] | 43 [62 28] | 39 [49 15] |
| **IC** | -58 [-53 -62] | -110 [32 -357] | -3.8 [-1.3 -7.6] | 38 [60 19] | 85 [174 18] |
| **PPGN** | -52 [-51 -53] | -39 [-4 -121] | -6.4 [-4.5 -8.5] | 66 [97 39] | 24 [50 8] |

*EUS: External urethral sphincter, IC: Ischiocavernosus, PPGN: Parasympathetic preganglionic neuron, RNSC: Region of negative slope conductance, AHP: Afterhyperpolarisation. Median [95% CI]*

### Somatic motoneurons receive recurrent synaptic inputs that are absent in parasympathetic preganglionic neurons

To investigate local synaptic organisation, we examined the distribution of En1+ and Calbindin+ synaptic terminals on retrogradely labelled perineal neurons in Ai34d × En1-Cre transgenic mice (**Figure 4A**). The colocalisation of these two markers identifies putative recurrent inhibitory inputs from Renshaw cells ^(26)^. Both EUS and IC motoneurons exhibited abundant En1+/Calbindin+ terminals (**Figure 4B**). In contrast, PPGN were devoid of these inputs, suggesting a distinct lack of recurrent inhibition in the autonomic control of the bladder.

**Figure 4.**
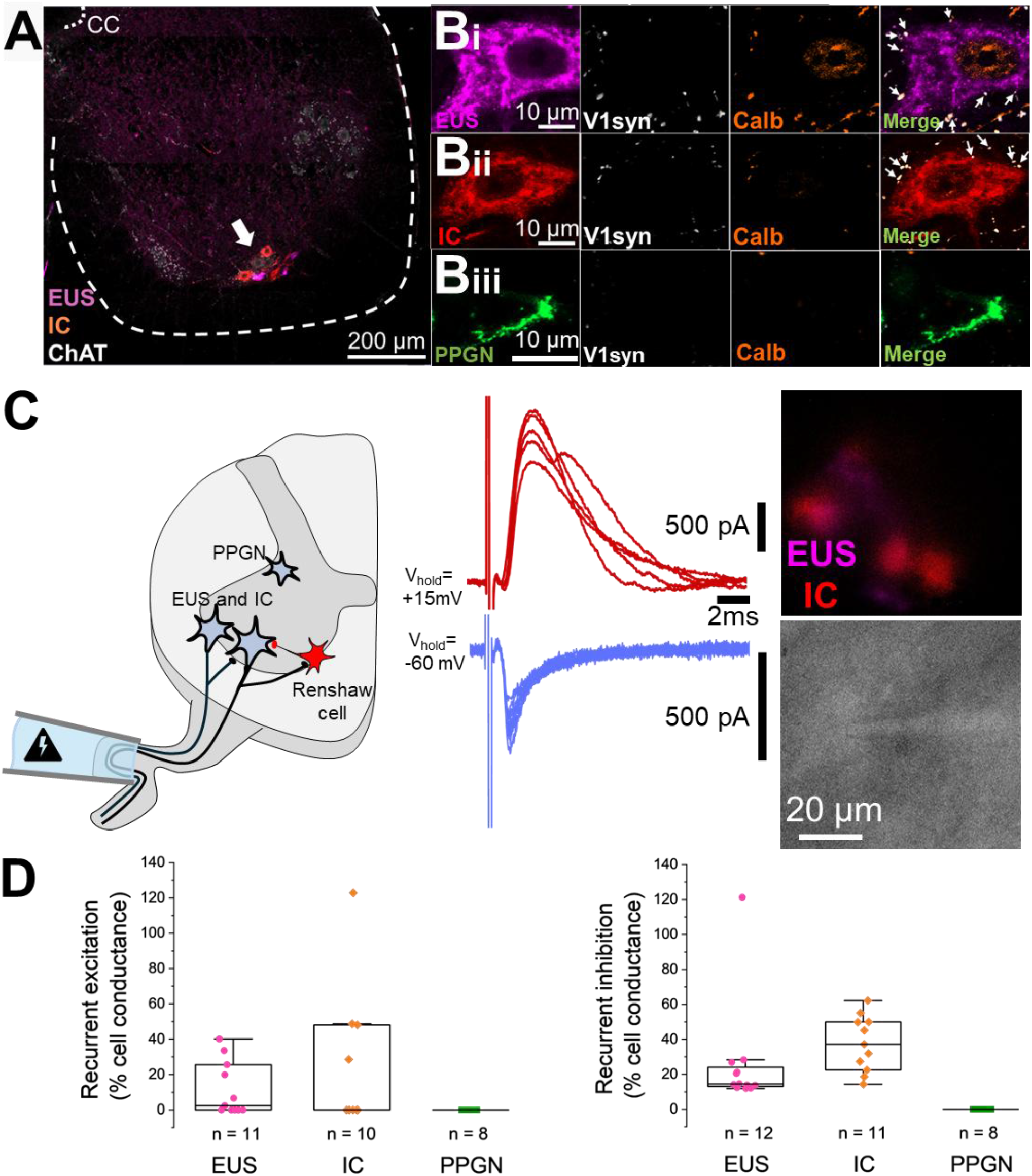
Anatomical and functional characterisation of recurrent inputs to autonomic parasympathetic preganglionic neurons (PPGN) and somatic motoneurons. **(A)** Image illustrating the retrograde tracing for external urethral sphincter (EUS) and ischiocavernosus (IC) motoneurons. **(B)** Immunolabelling of inhibitory inputs in the L6-S1 spinal cord. High-magnification insets show **(Bi)** EUS and **(Bii)** IC motoneurons receiving En1 (synaptophysin) and calbindin inputs, whereas **(Biii)** PPGN do not. Anatomical data obtained from N = 3 male Ai34d × En1-Cre mice (85– 95 days old). **(C)** Schematic of the experimental configuration for assessing recurrent circuits and representative voltage-clamp traces of responses evoked by ventral root stimulation. **(D)** Quantification of recurrent excitatory (left) and inhibitory (right) conductance scaled to cell conductance. Electrophysiological data collected from “n” number of neurons across 11 mice (10 males, 1 female; 42–111 days old).

To test whether these anatomical findings translate into function, we performed whole-cell patch-clamp recordings on retrogradely labelled perineal neurons. We assessed recurrent circuits in oblique slices via ventral root stimulation, which antidromically activates motor axons to recruit recurrent pathways (**Figure 4C**). Specifically, motor axons excite other motoneurons ^(27,28)^ and interneurons, including V3 ^(29)^, dI3 ^(30)^, and ventral spinocerebellar tract neurons ^(31)^ (recurrent excitation), as well as Renshaw cells (recurrent inhibition) ^(32–34)^. Ventral root stimulation elicited variable recurrent excitatory response (%cell responded, 54.5 for EUS, 40 for IC, 0 for PPGN). Also consistent with our anatomical data, ventral root stimulation evoked consistent recurrent inhibitory responses in all the somatic motoneurons. In the majority of recorded somatic motoneurons, recurrent inhibition and excitation were of comparable magnitude, although IC motoneurons received moderately stronger inhibition. In contrast, we detected no recurrent excitation or inhibition in any of the recorded PPGN neurons, confirming the lack of recurrent connectivity in this autonomic population (**Figure 4D**).

### Acute tibial nerve stimulation evokes short-latency synaptic potentials on sphincter motor units

Having mapped the local synaptic inputs to perineal neurons, we next investigated how peripheral nerve stimulation could impact the activity of EUS motor units. While chronic tibial nerve stimulation is an established therapy for overactive bladder ^(12,13)^, its underlying physiological mechanisms remain poorly understood. We hypothesise that acute tibial nerve stimulation elicits direct synaptic responses in EUS motor units via local spinal microcircuits.

To investigate the role of tibial nerve stimulation during EUS activity in urination, we employed a novel **‘pressure-clamp**’ technique using 3FR Fogarty catheters to maintain stable bladder distension, thereby ensuring constant, isometric-like EUS motor unit firing (**Figure 5A**). Under these conditions, unilateral tibial nerve electrical stimulation (4x the threshold of the toe twitch) evoked a distinct response on EUS motor unit activity. The stimulation elicited an early inhibitory response followed by a weak rebound excitation **(Figure 5B)**. We quantified the properties of the inhibitory response using a combination of PSTH and PSF analyses **(Figure 5C)**. Latency was defined as the time from stimulation onset to the first significant deviation in the PSTH-CUSUM ^(35)^. This value was relatively consistent across motor units (10.0 ± 3.8 ms, mean ± SD). The short latency of inhibition is consistent with local spinal circuits rather than supraspinal loops ^(36)^. Duration was measured as the interval between this onset and the return to baseline firing rates observed in the PSF-CUSUM ^(34,37–40)^. This was also consistent across conditions (50.0 ± 8.8 ms). Finally, the amplitude was assessed by the difference between the peak in discharge probability or rate and the trough following this peak, normalised to the number of stimuli delivered ^(41,42)^. Unlike the latency and duration, the amplitude derived from both PSTH-CUSUM (0.09 ± 0.07) and PSF-CUSUM (1.09 ± 0.96) showed considerable variability.

**Figure 5.**
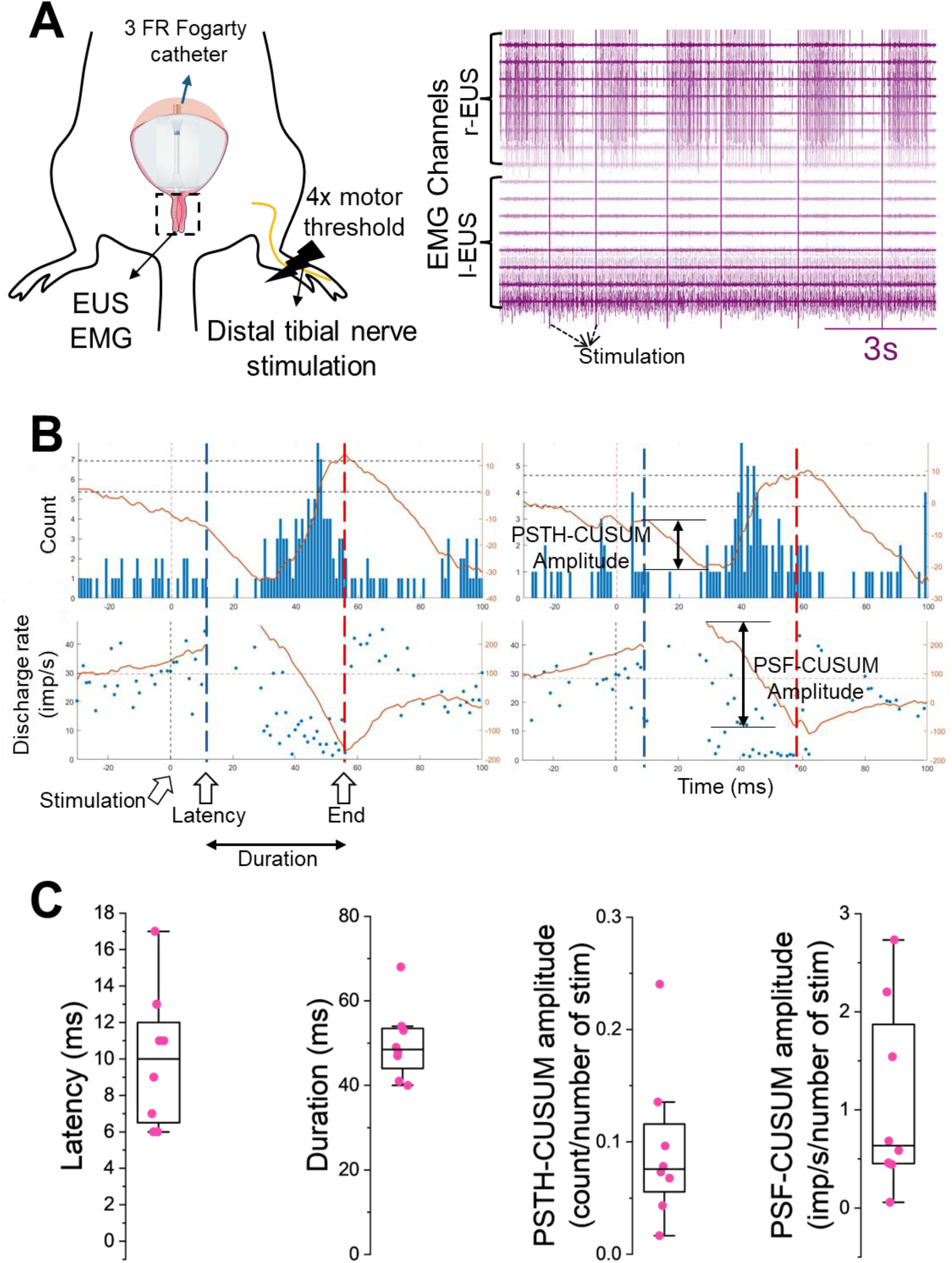
Modulation of external urethral sphincter (EUS) motor units by tibial nerve stimulation during bladder pressure-clamp. **(A)** Schematic illustrating the experimental protocol (left) and a representative raw myomatrix EMG trace of EUS motor unit activity during the pressure-clamp procedure (right). **(B)** Analysis of stimulus-evoked inhibitory response. Upper: Peristimulus time histogram (PSTH) showing the probability of motor unit discharge relative to tibial nerve stimulation. The red trace indicates the cumulative sum (CUSUM) used to detect significant deviations. Lower: Peristimulus frequencygram (PSF) displaying the instantaneous firing rates. Vertical dashed lines mark the onset latency (in PSTH) and the offset (in PSF) of the evoked response, and the estimated duration of the event. (**C**) Properties of inhibitory response including the latency, duration and amplitude (through PSF and PSTH CUSUM). Total of 8 motor units recorded from 2 male mice. Number of stimuli delivered to the tibial nerve was around 140 per motor unit.

## Discussion

The spinal control of urination requires coordination between local autonomic and somatic circuits yet accessing and characterising these networks in mature systems has been challenging. In this study, we developed a methodological framework, spanning *in vivo* HDEMG to decode motor unit activity in the EUS and IC, alongside *in vitro* whole-cell patch-clamp recordings of retrogradely labelled adult perineal neurons. Using this combined approach, we demonstrate that the distinct functional components of the spinal circuitry governing urination can be systematically investigated.

### Combining cystometry with intramuscular high-density EMG

Traditional methods for evaluating sphincter activity during cystometry typically rely on fine-wire EMG in animals ^(43,44)^, which capture a global, superimposed signal of overall muscle activation. While these conventional approaches are sufficient for identifying general reflex phase transitions (e.g., distinguishing storage from voiding), they fundamentally lack the spatial resolution required to reliably decompose the signal into individual motor unit action potentials. By integrating HDEMG with real-time cystometry, our methodology provides the spatiotemporal precision necessary to simultaneously track the activity of multiple individual motor units across complete urinary cycles and thus provide insight into spinal cord urinary circuit function. The ability to resolve single-unit activity *in vivo* is essential for decoding the neural drive to perineal muscles. Capturing these high-resolution recruitment and de-recruitment profiles is vital for future translational research, as it provides a precise baseline to identify how specific motor unit coordination and intermuscular synergies are altered in pathological conditions.

### Distinct biophysical properties of perineal neurons

To understand how motoneurons integrate inputs to generate these specific motor outputs, we investigated their intrinsic electrophysiological properties; however, performing whole-cell patch-clamp recordings on adult spinal tissue presents significant technical hurdles. The viability of the spinal cord slices typically declines after the first three postnatal weeks ^(45)^, making robust recordings from mature motoneurons notoriously difficult. However, the lower lumbar and sacral segments exhibit a relative resistance to hypoxia during the slicing procedure compared to cervical and upper lumbar regions, permitting viable recordings in adult tissue ^(46–49)^. Although even the small EUS could be injected for retrograde labelling of its MNs in male and female mice, the IC muscle is sexually dimorphic and significantly underdeveloped in females ^(50)^, so our IC dataset was necessarily restricted to male mice. Thus, recordings were from identified MNs in adult preparations, a technique that could increase our understanding of the normal urinary physiology as well as pathophysiology of disorders affecting lower urinary tract -which are more frequent in adults.

Our data highlights a fundamental biophysical divergence between the somatic (EUS and IC) and autonomic (PPGN) neuronal populations. While the majority of EUS motoneurons exhibit an immediate firing phenotype, PPGN and IC motoneurons predominantly display a delayed firing profile. Unlike previous observations in neonatal lumbar spinal cord, where immediate and delayed firing phenotypes reliably correlate with smaller and larger motoneuron sizes respectively ^(25)^, our adult recordings demonstrate that this relationship is not conserved. We observed that PPGN neurons are significantly smaller than EUS motoneurons, evidenced by their lower whole-cell capacitance (69 pF vs 167 pF), conductance (7 nS vs 29 nS) and rheobase (56 pA vs 398 pA), yet they predominantly exhibit the “fast-like” delayed firing pattern. Conversely, EUS motoneurons are larger but dominantly display the “slow-like” immediate firing phenotype. Additionally, PPGN neurons showed a much larger mAHP amplitude and slightly longer half-width, properties that likely reflect their overall slower discharge characteristics. This divergence could reflect the postnatal maturation and/or different functionality of pelvic motoneurons compared to the more commonly studied limb motor pools, challenging the direct extrapolation of classical developmental properties to the mature perineal motor system.

On the other hand, hyperpolarisation-activated currents, mediated by cation channels (HCN), play an important role in setting the resting membrane potential and regulating neuronal discharge ^(51)^. In our recordings, IC motoneurons exhibited uniquely large H-currents (~85% of cell conductance) compared to sphincter neurons. As the H-current is also known to regulate recruitment thresholds and action potential generation ^(52)^, this may underlie the distinct “burstlet” firing observed in a subset of IC motoneurons.

## Spinal circuits for perineal neurons

The output of motoneuron is governed by both their intrinsic membrane properties and synaptic inputs they receive. Anatomical evidence from our study revealed a differential organisation of motor inputs. Specifically, we observed abundant putative Renshaw cell inputs (En1+/Calbindin+) on EUS and IC motoneurons, suggesting strong recurrent inhibitory connectivity. The existence of recurrent inhibition within Onuf’s nucleus has been a subject of debate ^(53,54)^. Here, whole-cell recordings confirmed the recurrent motor connectivity observed in our anatomical data. Ventral root stimulation elicited robust recurrent inhibitory and excitatory postsynaptic currents in somatic motoneurons (EUS and IC), similar to the feedback loops described in limb motor control. In contrast, PPGN neurons were entirely devoid of recurrent inputs, reinforcing the functional segregation between the somatic and autonomic components of the pelvic floor. This distinct segregation is a critical feature to consider when investigating disease mechanisms or developing neuromodulation techniques, as these interventions may directly affect or rely on these recurrent motor circuits.

### Potential mechanisms of action for tibial nerve stimulation

Percutaneous tibial nerve stimulation is an established clinical therapy for alleviating symptoms of overactive bladder and provides a non-surgical alternative for sacral nerve stimulation ^(12,13)^. The stimulation is delivered via distal electrodes and effective when applied chronically over the course of 10-12 weeks (30 mins period every week) ^(55)^, yet its precise mechanism of action remains largely speculative. Current theories suggest that the therapeutic benefits arise from long-term post-stimulation inhibitory effects ^(14,15)^ or the modulation of µ, κ, or δ opioid receptors ^(56)^, which may alter the sensitivity of spinal reflex pathways over time. However, while previous studies have primarily focused on bladder activity, dysfunction in EUS motor control could also cause incontinence and aberrant intravesical pressure fluctuations. Consequently, we investigated the direct effect of tibial nerve stimulation on EUS motor unit activity.

To address this, our “pressure-clamp” method, combined with stimulus-triggered analysis ^(35,39–41)^ allowed us to isolate the direct, acute synaptic effects of tibial nerve stimulation on EUS motor units without the confounding variables of changing bladder volume/pressure. We demonstrate that acute tibial nerve stimulation evokes a distinct biphasic response: a robust, short-latency inhibition followed by a weaker rebound excitation. The latency of this inhibition (10.0 ± 3.8 ms) is too short to involve supraspinal loops, suggesting that tibial nerve stimulation acts through local spinal circuits.

The tibial nerve is a mixed nerve originating from the lumbosacral spinal cord, sharing close segmental proximity with the perineal motor pools. The stimulation parameters typically employed could also favour the recruitment of large-diameter alpha motor axons alongside sensory fibres ^(57–59)^. Thus, the electrical stimulation can generate antidromic volleys in tibial motor axons ^(60)^ that propagate to the lumbosacral spinal cord. Because recurrent motor collaterals are known to project across lumbosacral spinal segments ^(27)^, these antidromic volleys have the capacity to directly access the local perineal microcircuits, those involving Renshaw cells that we identified projecting to EUS and IC motoneurons. Although our present dataset could not prove a direct causal evidence, the temporal characteristics of the response, specifically the evoked inhibition duration (~50 ms), are similar to the duration of lumbar recurrent inhibition reported in mice ^(36)^. Thus, the modulation of recurrent inhibition could play an important role in the mechanisms of tibial nerve stimulation for treating overactive bladder.

## Limitations

There are several limitations of this study to consider. The reduced preparations used to record neuronal properties are nervous systems in which the circuits for micturition have by definition been interrupted, such that properties and post-synaptic currents may not reflect those in intact animals. In addition, both our cystometry and tibial nerve stimulation experiments were conducted in anaesthetised animals, which may suppress some synaptic pathways. Finally, if interested in human neural circuit control of urination, it should be noted that there are notable species differences in the voiding phase. In humans, voiding is characterised by the complete relaxation (silence) of the EUS, whereas the EUS in mice exhibit “burst” to expel urine.

## Conclusions

In summary, this study provides the foundation for characterising the spinal control of micturition. We developed electrophysiological and functional methods to characterise the intrinsic properties and segmental reflex circuits of perineal neurons. Furthermore, by studying single motor units, we defined the basic principles of motoneuron recruitment and de-recruitment across these pools. Collectively, these findings provide critical insights into the recurrent circuits and biophysical properties of the motoneurons innervating the perineal muscles, offering a mechanistic foundation for developing future targeted neuromodulation therapies for urinary dysfunction.

## Acknowledgement

MGO is supported by a Royal Society Newton International Fellowship NIF\R1\192316. MGO, RMB and MB are supported by a Brain Research UK grant PG23-100019. FN is supported by a Sir Henry Wellcome Postdoctoral Fellowship 221610/Z/20/Z. APV is supported by the European Union’s Horizon Europe research and innovation program under the Marie Skłodowska-Curie grant agreement no. 101151398. RMB and MB are funded by Wellcome Trust Discovery Award 227433/Z/23/Z, and RMB is supported by Brain Research UK. Myomatrix arrays were supplied by the Center for Advanced Motor BioEngineering and Research (CAMBER) which is supported by NIH grants U24NS126936 and R01NS109237 ^(16)^.

## Author contributions

Conceptualisation: MGO, RMB and MB, Writing-original draft: MGO, Writing-review and editing: MGO, FN, APV, KD, VB, RMB and MB, Investigation: MGO, FN, APV, KD and VB, Methodology: MGO, FN and APV, Data curation: MGO, FN, APV, KD and VB, Supervision: MGO, RMB and MB, Data analysis: MGO, FN and APV, Visualisation: MGO, FN and APV.

## Materials and Methods

### Animals

All animal experiments were conducted at University College London under the approval of the institutional ethical review committees. Procedures complied with the UK Animals (Scientific Procedures) Act 1986 and adhered to UK Home Office regulations (project licence PP2688499). We used both male and female mice with C57BL/6J background. The number of animals used per experimental protocol is reported in the Results section.

### In vivo HDEMG recording

#### Surgery and recording

Animals were initially anaesthetised with isoflurane (4% for induction, 2.5% for maintenance) and placed on a heating pad to maintain a skin temperature of ~31°C. At this stage, urethane (1.2 g/kg in a 1 mL volume PBS) was administered subcutaneously. Following abdominal shaving, a midline incision was made through the skin, peritoneum and abdominal musculature, and the surgical field was kept open using retractors (Surgical suite: SURGI-M, Tip: SURGI5017, Fixator: SURGI-5010-2, Kent Scientific, CT). The EUS and IC muscles were anatomically identified relative to the bladder and pubic symphysis (Figure 3A). Myomatrix arrays (RF-4x8-BVS-8, Camber, GA) were sutured to the surface of the target muscles using 8-0 Vicryl Rapide. Bilateral arrays (8 channels each) were implanted on the EUS, alongside a single array on the IC.

For cystometry, the apical surface of the bladder dome was punctured to insert flared polyethylene-10 (PE10) tubing, which was secured tightly with a 7-0 silk suture (18020-70, Interfocus, UK). Following a leak test via manual saline infusion, isoflurane delivery was terminated. After a 20-minute washout period, continuous cystometry was initiated via saline infusion at 25 μL/min ^(17)^ using a syringe pump (PHD Ultra, Harvard Apparatus, MA). Intravesical pressure was recorded in real-time using an APT300 force transducer and an HSE multi-sensor amplifier (73-5078; Harvard Apparatus, MA). Intramuscular electromyographic signals from the Myomatrix arrays were acquired using an RHD 32-channel headstage (C3324) connected to an RHD 512-channel Recording Controller (C3004; Intan, CA). Data were sampled at 20 kHz and band-pass filtered at 1-8000 Hz. The animal was grounded via the base of the tail.

#### Motor unit decomposition

To isolate individual motor unit spike trains from the monopolar HDEMG recordings, a validated blind source separation algorithm was applied ^(61,62)^. This approach resolves the convolutive mixing model of the EMG signal by calculating separation vectors (motor unit filters) to identify the original neural sources. Prior to decomposition, the raw signals were processed through a second-order Butterworth band-pass filter (100 to 5000 Hz), and channels demonstrating a poor signal-to-noise ratio were visually identified and excluded. Because of the long recordings, the motor unit filters were initially derived from a 30 s window selected during maximum muscle activation during the voiding phase. These established filters were then applied across the entire recording duration to track the identified MUs across all cycles. From the extracted motor units, only those exhibiting a pulse-to-noise ratio ≥ 28 dB were retained for further analysis ^(63)^. Finally, to calculate the onset time of the motor unit action potential relative to the algorithm’s identified discharge times, spike-triggered averaging was performed across all EMG channels ^(64)^. The specific channel displaying the maximum peak-to-peak amplitude was selected for this estimation. Using this channel, the motor unit action potential onset was defined at a threshold of 20% of the maximum motor unit action potential amplitude, and this onset was visually verified for every motor unit included in the final dataset.

#### Motor unit firing properties

All the individual motor unit features were computed as the average across all the voiding cycles per animal. Motor unit spike trains retained after decomposition were used to compute the instantaneous firing rate ^(65)^. The maximum firing rate was calculated as the average of the three maximum continuous instantaneous firing rates.

Motor unit recruitment and de-recruitment were characterised relative to bladder pressure. The recruitment threshold was defined as the bladder pressure at which the motor unit produced its first sustained firing. The de-recruitment threshold was similarly defined as the bladder pressure at which the motor unit ceased sustained firing during the descending phase of bladder pressure following voiding. Both thresholds were normalised to the peak bladder pressure recorded per animal. To assess the relationship between recruitment order and motor unit firing properties, linear regression analyses were performed between the recruitment and de-recruitment thresholds and the maximum firing rate for each animal.

Recruitment hysteresis was assessed for each motor unit by comparing the recruitment and de-recruitment thresholds. Motor units whose de-recruitment threshold exceeded their recruitment threshold (i.e. that continued firing at pressures below their recruitment level) were classified as exhibiting sustained firing. Conversely, motor units whose de-recruitment occurred at a higher bladder pressure than recruitment were classified as exhibiting non-sustained firing. To assess whether hysteresis varied across the motor pool, the relationship between the hysteresis and maximum firing rate was examined using linear regression models per animal.

#### Burst analysis during the voiding phase

During the voiding phase, EUS motor units transition from tonic to phasic bursting activity. Individual bursts were automatically identified from the EMG signals, and all bursting features were computed as the average across all voiding cycles per animal.

EMG signals were rectified, averaged across channels within the array, and band-pass filtered between 3 and 12 Hz to isolate the bursting activity. Burst frequency was estimated from the peak of the power spectral density computed using Welch’s method. Burst peaks were then identified from the filtered signal, and the onset and offset of each burst were determined using an adaptive amplitude threshold. For each identified burst, the following parameters were extracted: burst duration, defined as the interval between burst onset and offset; the number of motor unit firings contained within each burst, expressed as an as a proportion of the total number of firings across motor units; and bursting activity onset and offset thresholds, defined as the bladder pressure at which the first burst began and the last burst ended within the voiding phase, respectively. To assess whether specific motor unit subpopulations preferentially contribute to bursting, the number of spikes per burst was compared between earlier-recruited (higher firing rate) and later-recruited (lower firing rate) motor units using linear regression models per animal. Burst properties were reported as median values with 95% confidence intervals across animals.

#### Coherence analysis

Coherence analysis was employed to assess the degree of shared synaptic input both within the EUS motor pool (intramuscular coherence) and between the EUS and IC motoneuron pools (intermuscular coherence). All the coherence analyses were performed on concatenated recordings windows including the complete voiding cycles. Two complementary approaches were used for intramuscular coherence estimation. In the first approach, decomposed motor unit spike trains were used to generate cumulative spike trains. The identified motor units were randomly divided into two equally sized groups, and the spike trains within each group were summed to generate two cumulative spike trains ^(66)^. The coherence between the resulting spike trains was calculated using MATLAB’s Neurospec 2.11 toolbox. This procedure was repeated across 50 random permutations of motor unit groupings, and the resulting coherence spectra were averaged to obtain a stable estimate ^(67)^. In the second approach, the differential and rectified EMG signals from the two most spatially separated electrodes within the EUS array were used for coherence estimation using the same method. This approach enabled coherence estimation even when the number of reliably decomposed motor units was limited.

For intermuscular coherence between the EUS and IC, the EMG signals from each muscle were used since motor unit decomposition from the IC was not systematically available across all recordings. Coherence was computed between the rectified EMG from ipsilateral EUS and IC electrodes in one male mouse. The 95% confidence limit for significant coherence was calculated as 1 − (0.05)^(1/(L−1))^, where L denotes the length of segments used in the spectral estimation ^(68)^.

#### Tibial nerve stimulation

To evaluate reflex responses to tibial nerve stimulation, the continuous cystometry setup was modified into a “pressure-clamp” preparation. Instead of the PE10 tubing, a 3FR Fogarty catheter was inserted into the bladder to achieve controlled, static distension. The distension volume was calibrated to maintain a stable, continuous baseline of EUS motor unit activation. For peripheral nerve stimulation, two multi-stranded, coated stainless steel wires (25 μm strand diameter; A-M Systems, USA) were de-insulated at the tips to increase the active surface area, bent into a fish-hook shape for tissue anchoring and inserted subcutaneously adjacent to the medial malleolus using a 25G needle. The motor threshold was defined as the minimal current required to elicit visible toe movement. Stimuli (200 μs pulse width) were delivered at 4× threshold ^(15)^ with an inter-stimulus interval of 1–2 s, yielding 138 ± 64 stimuli per recording. During stimulation, continuous EUS activity was monitored; if the number of active motor units visibly declined, the intravesical pressure via the catheter was incrementally increased to restore and maintain a constant level of motor unit activation throughout the recording.

### In vitro electrophysiology

#### Retrograde labelling

To retrogradely label the motoneurons innervating the perineal muscles, we performed intramuscular injections of fluorescently conjugated Cholera Toxin subunit B (CTB) 3–7 days prior to electrophysiological recordings. Surgical depth of isoflurane anaesthesia (4% for induction, 2.5% for maintenance) was confirmed by the absence of foot and tail withdrawal reflexes. Following animal securement and the subcutaneous administration of buprenorphine, a midline abdominal incision was made to expose the EUS and IC muscles (Figure 3A). The surgical site was kept open using retractors (Surgical suite: SURGI-M, Tip: SURGI5017, Fixator: SURGI-5010-2, Kent Scientific, CT). We delivered CTB-Alexa Fluor 488 or CTB-Alexa Fluor 555 (0.2% wt/vol in PBS, Thermo Fisher) using a Hamilton syringe fitted with a pulled glass micropipette (20–40 μm tip diameter). For the EUS, bilateral injections were performed at both a proximal and a distal site (0.5 μL per site). For the IC, a single 0.5 μL injection was delivered bilaterally. To minimise leakage, the micropipette was left in place for 30s following each injection. The abdominal musculature (8-0 Vicryl Rapide, Ethicon Johnson & Johnson, UK), peritoneum (8-0 Vicryl Rapide, Ethicon Johnson & Johnson, UK), and skin (7-0 Vicryl Ethicon Johnson & Johnson, UK) were sequentially sutured. Mice recovered in a temperature-controlled heating chamber (30°C) before returning to their home cages.

#### Slice preparation

Mice were terminally anaesthetised via intraperitoneal injection of ketamine and xylazine (100 mg/kg and 10 mg/kg, respectively) prior to decapitation. The vertebral column was rapidly extracted and secured with insect pins ventral-side-up in ice-cold artificial cerebrospinal fluid (aCSF) containing (in mM): 113 NaCl, 3 KCl, 25 NaHCO_3_, 1 NaH_2_PO_4_, 2 CaCl_2_, 2 MgCl_2_, and 11 D-glucose, continuously aerated with 95% O_2_ and 5% CO_2_. The mid-lumbar to mid-sacral segments of the spinal cord were isolated. To generate oblique slices, the cord was glued at a 45° angle to a 6% agar block (supplemented with 0.1% methyl blue for visual contrast), with the ventral surface and intact ventral roots oriented toward the cutting blade. Sectioning was performed using a Leica VT1200 vibratome and the tissue was submerged in an ice-cold (~2°C) slicing solution consisting of (in mM): 130 K-gluconate, 15 KCl, 0.05 EGTA, 20 HEPES, 25 D-glucose, 3 kynurenic acid, 2 Na-pyruvate, 3 Myo-inositol, and 1 Na-L-ascorbate (pH 7.4 adjusted with NaOH). Oblique slices (350 μm thick) spanning the L6–S2 segments were collected, incubated in standard recording aCSF at 37°C for 30 min, and subsequently maintained at room temperature under continuous oxygenation (95% O_2_ / 5% CO_2_).

#### Identification of labelled neurons

Somatic motoneurons and PPGNs were differentiated based on their distinct anatomical distributions within the lumbosacral spinal cord; EUS and IC motoneurons reside in the ventrolateral Onuf’s nucleus, whereas PPGNs are located in the intermediolateral nucleus. Intramuscular EUS injections resulted in labelling of sacral PPGNs (presumably those innervating the internal urethral sphincter via proximal intramural ganglia, as reported in Fuller-Jackson, Osborne^(69)^). Consequently, anatomical location was used as the definitive criterion to distinguish EUS motoneurons from PPGNs when both were labelled with the same fluorophore. Distinct Alexa conjugates were used to separate EUS and IC motoneurons. Labelled neurons were visualised using a Nikon NI-FLTs epifluorescence microscope equipped with appropriate dichroic filters. Excitation was achieved by a 488-nm LED (for Alexa 488) and a white LED (for Alexa 555) (Opto LED, Cairns Instruments, UK), and images were captured via a Retiga XR CCD camera (QImaging, UK).

#### Whole-cell recording

Whole-cell patch-clamp recordings were obtained using an Axopatch 200B amplifier (Molecular Devices, Sunnyvale). Signals were low pass filtered at 5 kHz, digitised at 50 kHz via a Digidata 1440A A/D converter, and acquired using Clampex 10 software (Molecular Devices, Sunnyvale). Recording pipettes were fabricated from borosilicate glass capillaries (GC150F, Harvard Apparatus, UK) using a P-1000 Flaming-Brown puller (Sutter Instruments, CA) and polished with an MF2 microforge (Narishige). Final pipette resistances were 1–4 MΩ for somatic motoneurons and 3–6 MΩ for PPGNs.

The intracellular recording solution contained (in mM): 125 K-gluconate, 6 KCl, 10 HEPES, 0.1 EGTA, and 2 Mg-ATP, adjusted to pH 7.3 with KOH (osmolarity 290–310 mOsm). While we calculated a liquid junction potential of ~15 mV, the membrane potentials stated in the text and figures are reported as uncorrected values for simplicity.

Whole-cell capacitance and input resistance were calculated from the voltage response to a brief 200 ms, 50–100 pA current injection (current clamp). Resting membrane potential was measured prior to any current injection. To assess the firing phenotype and determine rheobase, 4-second depolarising current steps of increasing amplitude were applied until repetitive action potentials were elicited. The amplitude and half-width of the mAHP were quantified from the most negative trough to the pre-spike baseline using the initial, well-isolated spikes during the current step.

RNSCs were evoked in voltage-clamp mode via a slow depolarising ramp (10 mV/s from −70 to 0 mV). RNSC onset voltage and amplitude were extracted post-hoc from leak-subtracted traces ^(52,70,71)^. Hyperpolarisation-activated currents (H-currents) were measured during 1-second hyperpolarising voltage steps (from −60 to −110 mV, in 10 mV increments). The corresponding H-current conductance was determined from the difference between the instantaneous and steady-state currents and subsequently normalised to cell conductance.

To map recurrent synaptic inputs, ventral roots were stimulated via custom-fitted glass suction electrode using an isolated constant-current stimulator (DS3, Digitimer, UK). The motor threshold was defined as the minimum current required to reliably evoke a postsynaptic response in a somatic motoneuron. Subsequent test stimuli were delivered at 2–3 times this threshold. Excitatory and inhibitory postsynaptic currents (EPSCs and IPSCs) were recorded in voltage-clamp mode at holding potentials of −60 mV and +15 mV (before junction potential correction), respectively. Synaptic conductances were calculated directly from the peak current amplitudes, assuming a reversal potential of +15 mV for excitatory and −60 mV for inhibitory conductances (before junction potential correction).

### Immunohistochemistry

To visualise synaptic boutons and retrogradely labelled neurons, wild-type and transgenic En1 crossed with Ai34d (JAX:012570) mice were terminally anaesthetised and transcardially perfused with ice-cold phosphate-buffered saline (PBS), followed immediately by freshly prepared 4% paraformaldehyde (PFA) in PBS. The lumbosacral spinal cords were dissected, post-fixed with 4% PFA in PBS, and cryoprotected in 30% sucrose in PBS for 36 hours. Tissues were subsequently embedded in OCT compound (Tissue-Tek, 4583). Transverse sections (20 μm thickness) spanning the lumbosacral segments were obtained using a Leica CM3050 S cryostat.

Spinal cord sections were blocked for 30 minutes in a buffer comprising 0.2 M PBS, 0.2% Triton X-100 (Sigma, T9284), and 10% donkey serum (Sigma, D9663). Tissues were then incubated for 36 hours at 4°C with the following primary antibodies diluted in the blocking solution: goat anti-choline acetyltransferase (ChAT; 1:100, Millipore, AB144P) and rabbit anti-Calbindin (1:4000, Swant, CB38a). Following primary incubation, sections were washed and incubated overnight at 4°C in the same blocking buffer containing the following secondary antibodies: preabsorbed donkey anti-goat Alexa Fluor 405 (1:200, Abcam, ab175665) and donkey anti-rabbit Alexa Fluor 405 (1:500, Abcam, ab175649). Labelled sections were mounted with Mowiol (Sigma, 81381-250G) and cover slipped (VWR, #631-0147). Image acquisition was performed using a Zeiss LSM800 confocal microscope and ZEN Blue 2.3 software. Wide-field lumbosacral cords were captured using a 20× air objective (0.8 NA). High-resolution imaging of neurons and terminals was performed using a 63× oil-immersion objective.

### Disclosure and software

The bladder icons in Figures 1 and 2 were modified from Servier (smart.servier.com), licensed under CC-BY 3.0. The pubic bone image in Figure 2 was modified from DBCLS (togotv.dbcls.jp/en/pics.html), licensed under CC-BY 4.0. EMG signal acquisition was performed using an Intan RHD system (Intan Technologies, USA) and analysed with MATLAB R2025a (MathWorks, USA). Whole-cell patch-clamp recordings were acquired with Clampex 10 and subsequently analysed using Clampfit 11.4 (Molecular Devices, USA). Plots were generated using OriginPro 2024b (OriginLab Corporation, USA), Microsoft PowerPoint and Excel (Microsoft, USA) and MATLAB R2025a (MathWorks, USA).

